# Acetylation-dependent clustering of BRD2 instructs transcription dynamics

**DOI:** 10.1101/2025.09.22.677521

**Authors:** Niyazi Umut Erdogdu, Sukanya Guhathakurta, Maria Shvedunova, Ronald Oellers, Eric M. Patrick, Jose Alberto Morin Lantero, Janine Seyfferth, Ward Deboutte, Ibrahim I. Cissé, Asifa Akhtar

## Abstract

BET protein inhibitors represent one of the most prominent classes of epigenetic drugs. However, the specific contributions of individual BET family members to transcription regulation remain largely unknown. Here, by acutely co-depleting BET proteins, we uncover an essential role of BRD2 in maintaining RNA Polymerase II recruitment at promoters, which becomes especially critical in the absence of BRD4 or when pause release is inhibited. Combining rapid protein degradation, chemogenomics, and super-resolution microscopy, we show that H4 acetylation, particularly H4K16ac, serves as a template for BRD2 function, driving sub-diffraction BRD2 cluster formation at chromatin. Accordingly, depletion of BRD2 or the writer of H4K16ac, MOF decreases sub-diffraction RNA Polymerase II clusters, which are associated with transcription initiation. Thus, this study highlights the importance of acetylated chromatin in transcriptional coactivator organization and the distinct role of BRD2 in transcription initiation dynamics.

## MAIN TEXT

Bromodomain (BD) and extra-terminal (ET) domain (BET) family proteins, consisting of BRD2, BRD3, BRD4 and BRDT, are widely acknowledged as major transcriptional regulators and characterized by their signature structural organization as two N-terminal tandem BDs followed by an ET domain ^1^. Over the years, BET proteins have gained significant attention for their role in cancer biology, as they are pivotal activators of oncogenic networks in various tumors. The development of BET inhibitors, which disrupt the bromodomain-dependent chromatin binding of BET proteins, provided a promising tool for the suppression of oncogenic pathways ^2^.

Despite years of research on these proteins, there is only consensus on the role of BRD4 in the regulation of transcription elongation: BRD4 promotes the transition of proximally paused RNA Pol II into productive elongation by forming a catalytically active BRD4-pTEFb and facilitating the formation of functional elongation complexes ^3,4^. On the contrary, the transcriptional regulatory roles of BRD2 and BRD3 remain less well defined. It has been previously shown that both BRD2 and BRD3 enabled RNA Pol II to transcribe through nucleosomes containing specific histone H4 modifications such as H4K5ac and H412ac in a defined *in vitro* chromatin transcription system ^5^. However, their rapid depletion in cells did not lead to any major changes in global transcription elongation suggesting that these proteins might execute more nuanced regulatory functions in transcription, the molecular mechanisms of which are not fully characterized ^4^.

## H4-ac mediated by MOF is required for the recruitment of BRD2 to chromatin

It has been recently shown that H4 acetylation strongly recruits BET proteins to the chromatin via their bromodomains *in vitro* ^7^. Therefore, in this study, to understand the role of H4 acetylation in the control of transcription dynamics, we decided to deplete the MYST family lysine acetyltransferase KAT8/MOF, which primarily catalyses H4K16 *in vivo* and has been shown to additionally acetylate H4K5, H4K8, and H4K12 *in vitro* ^8,9^. Since the long duration of typical perturbation methods such as RNA interference (RNAi) and inducible Cre- or CRISPR-Cas9-mediated knockout can obscure the direct consequences of target depletion, a rapid degradation system for MOF was required to capture the dynamic changes in chromatin and transcription. To this end, we first generated an mESC line which stably expresses the E3 Ub ligase Tir1 from *O. sativa* from a cassette introduced into the *Rosa26* locus using TALEN (Tir1^+^ WT26 mESCs). Then, we endogenously tagged MOF with V5 and mAID tags in Tir1^+^ WT26 mESCs using CRISPR-Cas9 (Fig. 1a). Auxin treatment of MOF degron mESCs resulted in near-complete proteasomal degradation of MOF within 45 minutes and could be reversed by washing off auxin (Fig. 1b and Extended Data Fig. 1a, b). Bulk levels of H4K16ac were reduced only after 3 hours of auxin treatment, indicating that the turnover of this histone mark requires at least a few hours in the absence of new catalysis (Fig. 1b). The robustness of the degron system was further validated by ChIP-Seq showing near complete loss of enrichment of MOF at its chromatin binding sites after 3 hours of auxin treatment (Fig. 1c). In line with the literature, rapid depletion of MOF led to severe proliferation defects and cell death within a time-course of 96 h (Extended Data Fig. 1c, d) ^10,11^.

**Fig. 1:**
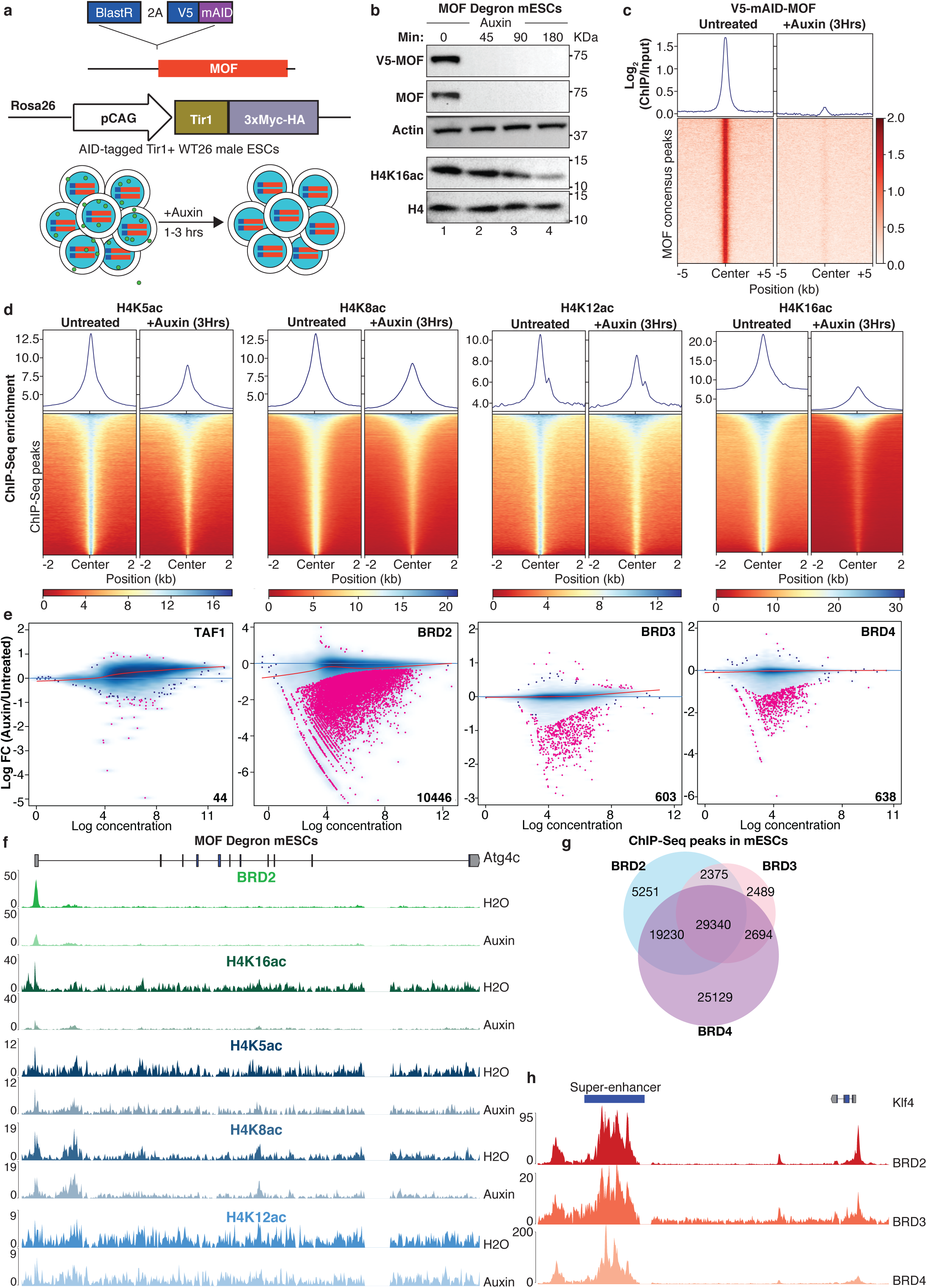
MOF-mediated H4 acetylation is required for the recruitment of BRD2 to the chromatin. **a**, Schematic showing the genome editing strategy to generate an auxin-inducible degron line for MOF. **b**, WB showing rapid degradation of MOF in MOF degron mESCs upon auxin treatment for indicated time points. Actin serves as a loading control. **c**, Metagene plot and heatmap of input-normalized ChIP-Seq signal for MOF from MOF degron mESCs upon auxin treatment for 3 h. The signal was plotted over MOF consensus peaks and MOF was immunoprecipitated using V5 antibody. **d**, Metagene plot and heatmap of spike in-normalized native ChIP-Seq signal for H4K5ac, H4K8ac, H4K12ac and H4K16ac over their respective peaks upon auxin treatment of MOF degron mESCs for 3 h. **e**, MA plots of differential chromatin enrichment analysis for TAF1, BRD2, BRD3 and BRD4 upon auxin treatment of MOF degron mESCs for 3 h. Pink dots depict differentially enriched peaks with FDR< 0.05. **f**, Genome snapshot of the MOF-sensitive BRD2 target gene *Atg4c* showing H4K5ac, H4K8ac, H4K12ac, H4K16ac and BRD2 ChIP-Seq signals upon auxin treatment of MOF degron mESCs for 3 h. **g**, Venn diagram depicting the distinct and overlapping peaks of BRD2, BRD3 and BRD4 ChIP-Seq peaks in mESCs. **h**, Example genome snapshot of BRD2, BRD3 and BRD4 ChIP-Seq signals over the super-enhancer region near the *Klf4* gene.

Taking advantage of the rapid degradation system, we next performed time-course total RNA-Seq experiments in order to determine the earliest time point of auxin treatment which generates a strong and direct transcriptional response to MOF depletion without induction of secondary effects. Volcano plots obtained from DESeq analysis indicated an increase of differentially expressed genes with increasing treatment time. The earliest time frame for auxin treatment by which we could observe a prominent transcriptional response was 3 h (Extended Data Fig. 1e). Considering that the onset of a bulk-level decrease in H4K16ac levels also occurred at 3 h of auxin treatment (Fig. 1b), we decided on 3 hours of auxin treatment for our experiments.

In order to investigate the direct contribution of MOF to H4 acetylation, we performed native ChIP-Seq experiments for the four H4 tail acetylation marks upon 3 h auxin treatment of MOF degron mESCs. Loss of MOF led to overall reduced levels of H4K5ac, H4K8ac, H4K12ac and H4K16ac, with the most severe effect on H4K16ac (Fig. 1d). 65% of all H4K16ac ChIP-Seq peaks were associated with chromatin features related to active transcription such as transcription elongation, active promoters and enhancers (Extended Data Fig. 1f-h).

The prevailing view in the literature is that histone acetylation is able to exert an influence on transcription through the binding of proteins with bromodomains which in turn recruit transcriptional cofactors that alter RNA Pol II activity ^12^. In particular, H4 acetylation has been implicated in the recruitment of BET proteins to the chromatin (Extended Data Fig. 1i) ^5,7,13,14^. Additionally, the TAF1 subunit of the general transcription factor TFIID has been shown to bind H4-acetylated peptides with high affinity and has been recently proposed to be recruited to transcription start sites by MOF-mediated H4 acetylation ^15,16^. Differential chromatin occupancy analysis for TAF1 and BET BRD proteins upon acute MOF depletion for 3 h revealed only minor changes in TAF1, BRD3 and BRD4 binding (Fig. 1e). However, the genome-wide chromatin occupancy of BRD2 was severely altered by MOF depletion as evident by 10,446 BRD2 binding sites showing loss of enrichment (Fig. 1e and Extended Data Fig. 1j). On MOF-sensitive BRD2 binding sites, there was a reduction of H4K5ac, H4K8ac, H4K16ac and to a lesser extent H4K12ac. However, the most prominent changes were again observed in H4K16ac (Fig. 1f and Extended Data Fig. 1k). Interestingly, there was a significant overlap in the overall chromatin binding sites of the BET BRD proteins in mESCs (Fig. 1g, h). However, we found that BRD3 and BRD4 remain bound near MOF-sensitive BRD2 peaks, indicating depletion of MOF and the reduced H4-ac selectively interferes with BRD2 chromatin binding dynamics (Extended Data Fig. 2a, b).

## Co-depletion of BET BRD proteins identifies their distinct roles in the regulation of nascent transcription

MOF-sensitive BRD2 peaks largely overlap with chromatin regions associated with active transcription (Fig. 2a). Given the limited literature on the role of BRD2 in transcription regulation, we were intrigued to investigate the functional significance of the H4 acetylation-BRD2 relationship in the regulation of transcription dynamics. To this end, we generated an auxin-inducible degron cell line for BRD2 by endogenously tagging it with an mAID tag using CRISPR-Cas9 in Tir1^+^ WT26 mESCs (Fig. 2b). Upon addition of auxin, near-complete proteasomal degradation of BRD2 was reached within 45 minutes (Fig. 2c). Furthermore, we could validate the robustness of the degron system by demonstrating the loss of BRD2 ChIP-Seq peaks upon auxin treatment for 3 hours (Extended Data Fig. 2c). We performed transient transcriptome sequencing (TT-Seq) experiments in BRD2 degron mESCs to quantitatively interrogate the function of BRD2 in RNA synthesis ^17^. In general, it was observed that nascent RNA enrichment in this experiment led to a better coverage throughout the gene body including introns and thus allowed a more sensitive transcriptional analysis (Extended Data Fig. 2d). In line with the literature ^4,18^, TT-Seq experiments indicated that rapid depletion of BRD2 alone did not affect RNA synthesis globally, but rather led to locus-specific mild transcriptional changes (Fig. 2d and Extended Data Fig. 2e).

**Fig. 2:**
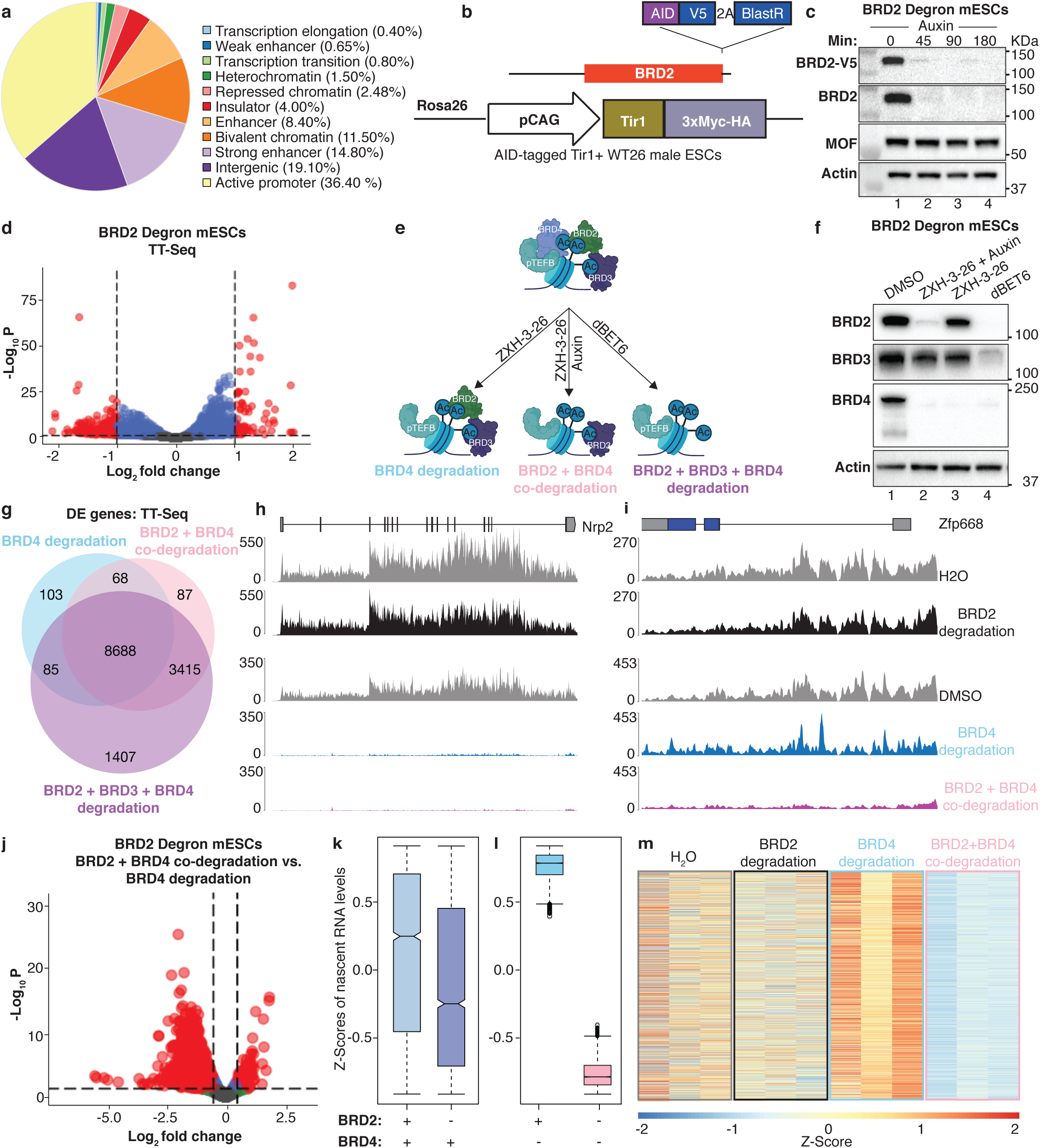
Acute co-depletion of BET proteins identifies their distinct roles in promoting RNA synthesis. **a**, Pie chart depicting the genome-wide distribution of MOF-sensitive BRD2 peaks sorted by chromatin features obtained from chromHMM analysis. **b**, Schematic showing the genome editing strategy to generate an auxin-inducible degron line for BRD2. **c**, WB showing the rapid degradation of BRD2 in BRD2 degron mESCs upon auxin treatment for different time points. Actin serves as a loading control. **d**, Volcano plot showing differentially expressed genes (red) identified by TT-Seq from BRD2 degron mESCs treated for 3 h with auxin. The dashed lines mark the log2FC-cutoff of 1 and adjusted p-value cutoff of 0.05. **e**, Schematic depicting the experimental strategy for co-depletion of BET BRD proteins. **f**, WB showing rapid degradation of BET BRD proteins upon combined treatments as described in (**e**). **g**, Venn diagram showing the overlap of differentially expressed genes (log2FC< -1 & p-adj< 0.05) detected in TT-Seq experiments upon co-depletion of BET BRD proteins. **h, i**, Genome snapshots of nascent RNA signals for representative genes *Nrp2* (**h**) and *Zfp668* (**i**) upon auxin treatment of BRD2 degron mESCs (upper two tracks) and combined depletion of BRD2 and BRD4 (lower three tracks). **j**, Volcano plot showing differentially expressed genes identified by TT-Seq (red) upon 3 h auxin treatment of BRD4-depleted BRD2 degron mESCs (genes deregulated upon combined BRD2 and BRD4 depletion compared to BRD4 depletion alone). The dashed lines mark the log2FC-cutoff of 0.5 (vertical line) and adjusted-p value cutoff of 0.05 (horizontal lines). **k, l**, Boxplots depicting the z-scores for nascent RNA levels of the significantly downregulated genes scored in **(j)** upon depletion of BRD2 alone **(k)** or upon co-depletion of BRD2 and BRD4 **(l)**. **m**, Heatmaps showing Z-score nascent expression of the significantly downregulated genes from **(j)** upon depletion of BRD2 or BRD4 alone and co-depletion of BRD2 and BRD4.

Given the fact that BET BRD proteins frequently co-occupy the same chromatin binding sites, we argued that they might play distinct, but collaborative roles in the regulation of nascent transcription. Therefore, we decided to perform co-depletion experiments of BET BRD proteins combined with transient transcriptome sequencing (TT-Seq) to unravel their functional organization in promoting RNA synthesis. To this end, we combined auxin treatment in BRD2 degron mESCs with the BRD4-specific PROTAC reagent ZXH-3-26 to co-deplete BRD2 and BRD4 ^19^. We have also independently treated the cells with dBET6, a PROTAC reagent inducing the proteasomal degradation of all three BET BRD proteins (Fig. 2e). By western blot, we could confirm the specificity of individual reagents and their combinations to degrade the target BET BRD proteins (Fig. 2f).

Unlike BRD2 depletion alone, depletion of BRD4 alone or all BET BRD proteins led to a severe defect in RNA synthesis with an overlap of 8,763 significantly downregulated transcripts (Log2FoldChange<-1 & adjusted p-value<0.05) (Fig. 2g, h and Extended Data Fig. 2f). Interestingly, there were an additional 3,415 transcripts only significantly downregulated upon the combined depletion of BRD2 and BRD4 or all three BRD proteins when compared to depleting BRD4 alone (Fig. 2g, i). Furthermore, the presence of 1407 transcripts that were significantly downregulated only upon depletion of all 3 BET BRD proteins, but not upon co-depletion of BRD2 and BRD4 suggest that the role of BRD3 in promoting RNA synthesis is possibly also not fully redundant with BRD2 or BRD4 (Fig. 2g).

PCA analysis revealed that combined depletion of BRD2 and BRD4 resulted in transcriptional profiles which are more similar to triple BET BRD depletion than those in which only BRD4 was depleted (Extended Data Fig. 2g). Indeed, a direct DESeq comparison identified 5,159 transcripts that were more severely downregulated upon co-depletion of BRD2 and BRD4 when compared to the BRD4 degradation alone (Fig. 2j). We also found that the presence of BRD2 becomes more crucial for sustaining RNA synthesis for these 5,159 transcripts in the absence of BRD4 as revealed by the increased magnitude of transcriptional downregulation upon BRD2 loss in cells lacking BRD4 compared to cells retaining BRD4 (Fig. 2k-m). By mathematical modeling of total RNA and nascent RNA levels, we could perform transcriptional rate calculations and similarly show that the impact of BRD2 loss on RNA synthesis rates for these transcripts was more severe in the absence of BRD4 (Extended Data Fig. 2h, i). Collectively, these findings indicated that BRD2 provides a support function to prevent a transcriptional collapse in the absence of BRD4.

## BRD2 maintains the chromatin occupancy of RNA Pol II in the absence of BRD4

To further dissect the functional divergence of BET BRD proteins in transcription regulation, we have also performed RNA Pol II ChIP-Seq experiments with BRD2 degron mESCs upon single or combined depletions of BRD2, BRD3 and BRD4. In line with the transcriptional profile we observed, globally, depletion of BRD2 alone resulted in a mild, but significant decrease in promoter RNA Pol II occupancy without any significant change in gene body RNA Pol II levels (Fig. 3a ,b). In agreement with the literature ^3,4^, reduced RNA synthesis upon acute depletion of BRD4 was accompanied by an increased enrichment of RNA Pol II at the promoters, reminiscent of defects associated with the release of paused RNA Pol II (Fig. 3c, d). Interestingly, despite the dramatic reductions in gene body-associated RNA Pol II levels and RNA synthesis rates upon combined depletion of all 3 BET BRD proteins, we did not observe an increase in promoter RNA Pol II occupancy. Moreover, co-depletion of BRD2 with BRD4 was sufficient to reverse the accumulation of paused RNA Pol II at the promoters induced by the depletion of BRD4 alone, suggesting that BRD2 potentially acts upstream of BRD4 in the control of RNA Pol II progression (Fig. 3c, d). In line with this finding, by performing mass spectrometry after immunoprecipitation of endogenous BRD2 in mESCs, we found that the strongest interactors of BRD2 are components of basal transcription machinery. Particularly, multiple subunits of the TFIID complex were enriched upon BRD2 pull-down implying that BRD2 might contribute to the regulation of transcription initiation (Fig. 3e-g, Supplementary Table 1).

**Fig. 3:**
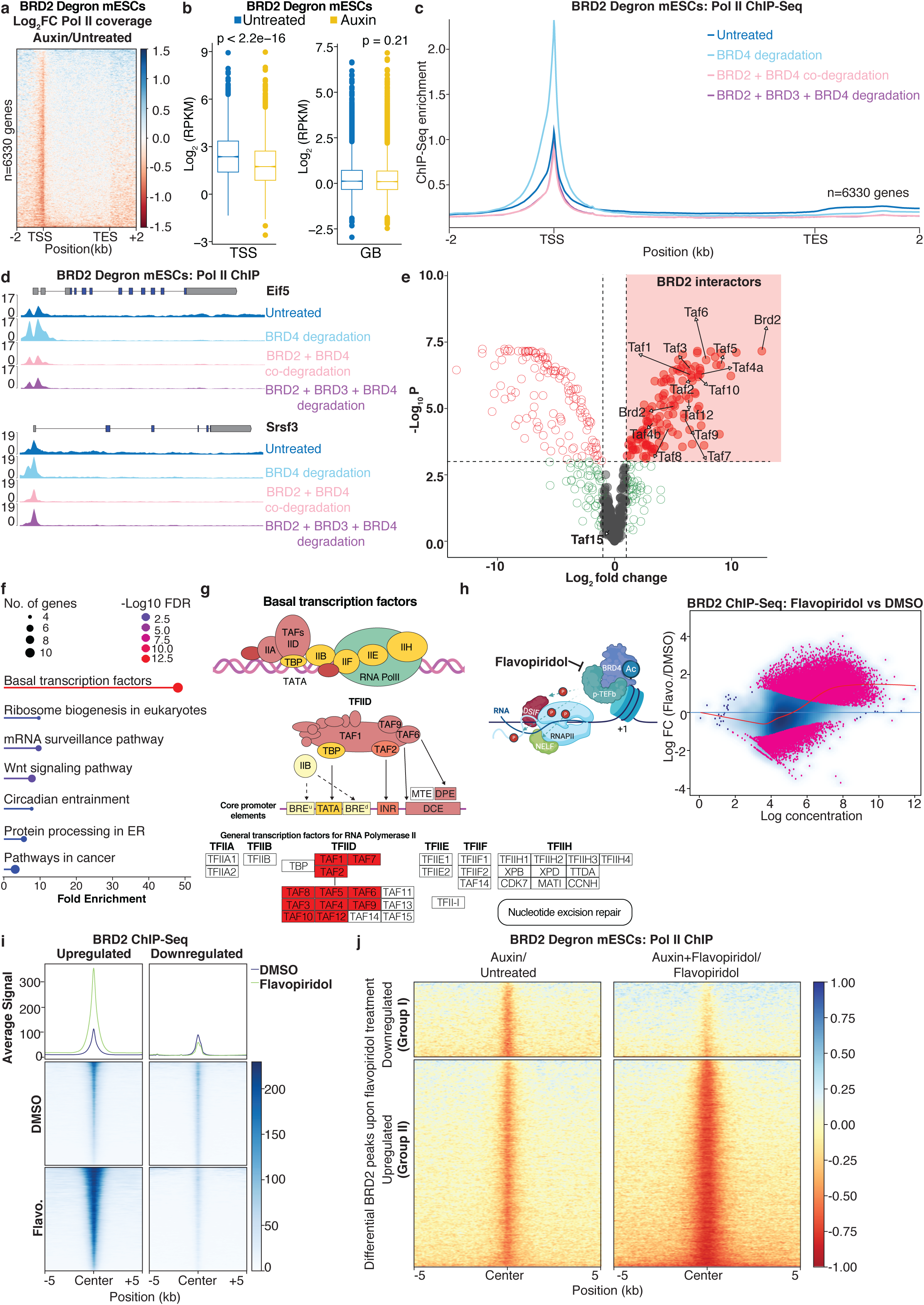
BRD2 regulates the chromatin occupancy of promoter-associated RNA Pol II. **a**, ChIP-Seq analysis depicting the Log2FC of RNA Pol II coverage upon auxin treatment of BRD2 degron mESCs for 3 h. **b**, Boxplots depicting RPKM-normalized RNA Pol II coverages over transcriptional start sites (TSSs) and gene bodies (GBs) upon auxin treatment of BRD2 degron mESCs for 3 h. **c**, Metagene plot showing RPM-normalized RNA Pol II ChIP-Seq signal upon co-depletion of BET BRD proteins in BRD2 degron mESCs. **d**, Genome snapshots depicting the changes in RNA Pol II ChIP-Seq signal on the representative genes *Eif5* (top) and *Srsf3* (bottom). **e**, Volcano plot depicting significant BRD2 interactors (log2FC>1 and FDR< 0.001) identified by mass spectrometry in comparison to the IgG negative control. **f**, GO term analysis for the significant BRD2 interactors. **g**, KEGG graph depicting the ‘basal transcription factors’ category with significant BRD2 interactors marked in red. **h**, Schematic describing the experimental strategy to study the function of BRD2 in relation to the dynamics of paused RNA Pol II using flavopiridol-mediated inhibition of pTEFb catalytic activity **(left)** and MA plots of differential chromatin enrichment analysis for BRD2 upon flavopiridol treatment of BRD2 degron mESCs for 1 h (no addition of auxin). Pink dots depict differentially enriched peaks with FDR< 0.05 **(right)**. **i**, Metagene plot and heatmap of ChIP-Seq signal for BRD2 in BRD2 degron mESCs upon flavopiridol treatment for 1 h (no addition of auxin). DMSO treatment served as a control and the signal was plotted over the differential BRD2 ChIP-seq peaks obtained from **(h, right)**. **j**, ChIP-Seq analysis depicting the log2FC of RNA Pol II coverage upon rapid BRD2 depletion with (right) and without (left) flavopiridol treatment. The signal was plotted over the differential BRD2 ChIP-seq peaks obtained from **(h)**.

Given the more prominent role of BRD2 in the control of RNA Pol II progression in the absence of BRD4, we next asked whether there might be a dynamic interplay between BRD2 and paused RNA Pol II. To this end, we inhibited RNA Pol II pause release using the pTEFb inhibitor flavopiridol in order to induce genome-wide RNA Pol II pausing in an independent manner from BRD4 depletion (Fig. 3h, left). Inhibition of pause release resulted in a massive accumulation of BRD2 on chromatin (Fig. 3h, right and 3i).

This unexpected finding prompted us to interrogate the impact of BRD2 depletion on RNA Pol II chromatin occupancy in relation to increased BRD2 enrichment upon pause release inhibition. To this end, we analyzed differential RNA Pol II coverage around the BRD2 peaks which either reduced (Group I peaks) or gained (Group II peaks) enrichment upon pause release inhibition (Fig. 3i). Similar to our previous observation (Fig. 3a), BRD2 depletion resulted in a mild reduction in RNA Pol II chromatin binding under steady state conditions in both groups (Fig. 3j, left). However, upon inhibition of pause release, BRD2 loss led to a more pronounced reduction in RNA Pol II occupancy only around Group II peaks (Fig. 3j, bottom), but not around Group I peaks (Fig. 3j, top). These findings imply a novel function of BRD2 in maintaining the chromatin occupancy of promoter-proximal RNA Pol II.

## Differential clustering properties of BRD2 and BRD4

Intrigued by the significant increase in BRD2 chromatin occupancy upon flavopiridol treatment, we wanted to follow the changes in the dynamics of BRD2 at single-cell resolution in this process. Hence, we endogenously tagged BRD2 at its C-terminus with a monomeric EGFP tag using CRISPR-Cas9. Under steady-state conditions, BRD2 showed a relatively homogenous distribution in the nucleus. However, upon addition of flavopiridol, BRD2 appeared in dynamic foci with a continuous dissolving and reappearance cycle (Fig. 4a and Supplementary Movie 1). We could demonstrate the dynamic nature of the BRD2 foci that form upon flavopiridol treatment using FRAP: After photobleaching, BRD2-mEGFP foci recovered fluorescence within 30 seconds (Fig. 4b). We could also quantify the emergence of the flavopiridol-induced BRD2 foci by performing k-means clustering analysis of the BRD2-mEGFP signal (Fig. 4c), which corroborates the increased BRD2 chromatin binding observed following flavopiridol treatment in our ChIP-Seq results (Fig. 3h, i). In order to confirm that BRD2 foci form due to its accumulation on chromatin upon pause-release inhibition, we inhibited its association with acetylated histones using JQ1. Indeed, low-dose JQ1 treatment diminished BRD2 foci induced by pause-release inhibition implying that acetylated chromatin template is a prerequisite for the spatial organization of BRD2 upon inhibition of pause-release (Fig. 4d).

**Fig. 4:**
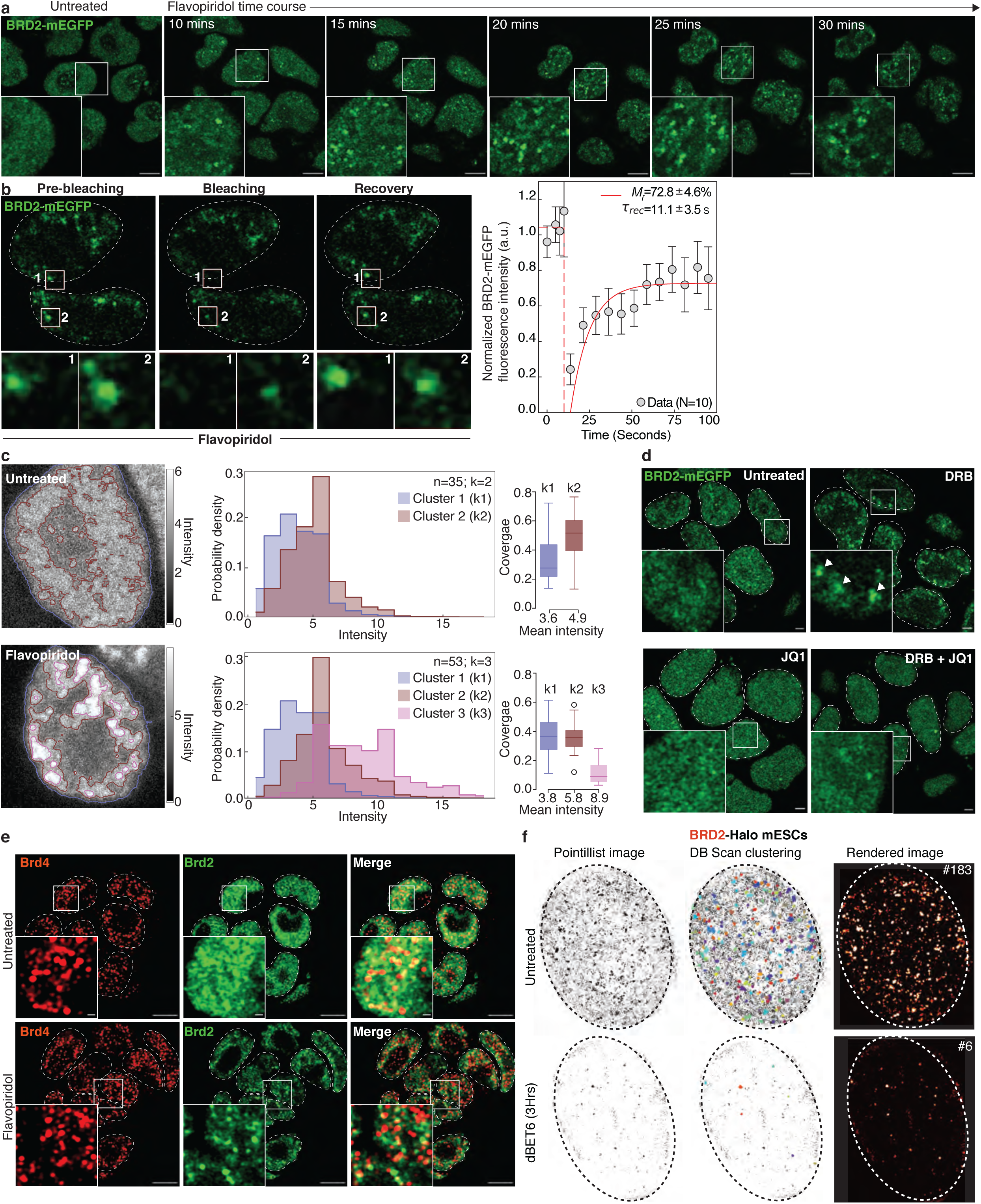
BRD2 re-organizes into dynamic foci upon inhibition of pause release. a, Time-lapse live-cell fluorescence microscopy images of BRD2-mEGFP signal in untreated and flavopiridol-treated (for the indicated time points) mESCs. Scale bar of images: 5 μm. **b**, Live-cell fluorescence microscopy images of flavopiridol-treated (30 mins) BRD2-mEGFP mESCs and relative quantification of fluorescence recovery kinetics of BRD2 clusters following bleaching. The FRAP curve shows mean 土 standard error across 10 independent clusters for individual time points. Scale bar of images: 5 μm. **c**, K-means classification of pixel intensities from live-cell images of cells treated with flavopiridol for 30 minutes. A contour plot showing each of the clusters detected (left), a histogram depicting probability densities of pixel intensities in each cluster detected (middle) and a box plot depicting nuclear coverage of each cluster detected (right) are shown. **d**, Live-cell fluorescence microscopy images of BRD2-mEGFP cells upon single or combined treatments with DRB and JQ1 for 1 h. Scale bar: 5 µm. **e**, Immunofluorescence microscopy images of untreated and 1 h flavopiridol-treated BRD2 degron mESCs stained using antibodies against BRD4 (red) and V5 (green, for BRD2-mAID-V5). **f**, Example of a super-resolved live-cell image of BRD2-Halo in mESCs. Two-dimensional super-resolution reconstruction reveals the presence of BRD2 clusters which disappear upon dBET6-mediated degradation of BRD2. The numbers on top right of the rendered images indicate the number of BRD2 clusters in the representative cells detected by DBSCAN.

Since BRD4 has been previously described to form stable clusters in mESCs, we next asked whether BRD2 foci colocalize with BRD4 clusters ^20^. To this end, we performed immunofluorescence experiments upon treatment of the cells with flavopiridol: While BRD4 clusters are already present under steady-state conditions, BRD2 foci appeared only upon flavopiridol treatment. BRD2 foci and BRD4 clusters only minimally overlap in flavopiridol-treated cells, implying distinct features in the nuclear dynamics of these two proteins (Fig. 4e).

The intrinsically disordered region (IDR) of BRD4 is larger in comparison to that of BRD2 despite their overall similar domain organization (Extended Data Fig. 3a, b). Nevertheless, the IDR of BRD2 was also able to phase-separate *in vitro* and these *in vitro* condensates were sensitive to both ATP and RNA (Extended Data Fig. 3c-e). Therefore, we hypothesized that BRD2 itself might also be capable of forming protein clusters that are shorter-lived in comparison to stable BRD4 condensates and which we therefore are not able capture with confocal microscopy. To test this hypothesis, we have endogenously tagged the 3’ end of the *Brd2* CDS with a Halo tag using CRISPR-Cas9 and observed the spatial dynamics of BRD2 using live-cell super-resolution microscopy. To perform time-correlated photoactivation localization microscopy (tcPALM), we have labeled BRD2-Halo with PA-JF646 in live mESCs. In line with our hypothesis, tcPALM measurements revealed the existence of transient BRD2 clusters which disappeared upon dBET6 treatment, confirming the specificity of the signal. BRD2 clusters exhibited an average size of ca. 250 nm and there were on average ca. 200 BRD2 clusters per cell (Fig. 4f and Extended Data Fig. 3f).

## H4-acetylation mediated by MOF is crucial for the regulation of BRD2 transient clusters and transcription dynamics

In order to investigate whether BRD2 clustering is crucial for its function in transcription regulation, we have generated BRD2 degron mESCs which express BRD2 variants that differ in their IDR under the control of a doxycycline-inducible promoter (Fig. 5a, b). We performed RT-qPCR for several target genes that are responsive to the co-depletion of BRD2 and BRD4 (Fig. 2j). While overall exogenous expression of WT-BRD2 could rescue the expression of target genes upon rapid co-depletion of endogenous BRD4 and BRD2, the BRD2-IDRdel variant failed to do so. Expression of BRD2-IDR^BRD^^4^ could largely rescue the expression of target genes, suggesting that BRD2 clustering is critical for its function in transcription regulation (Fig. 5c).

**Fig. 5:**
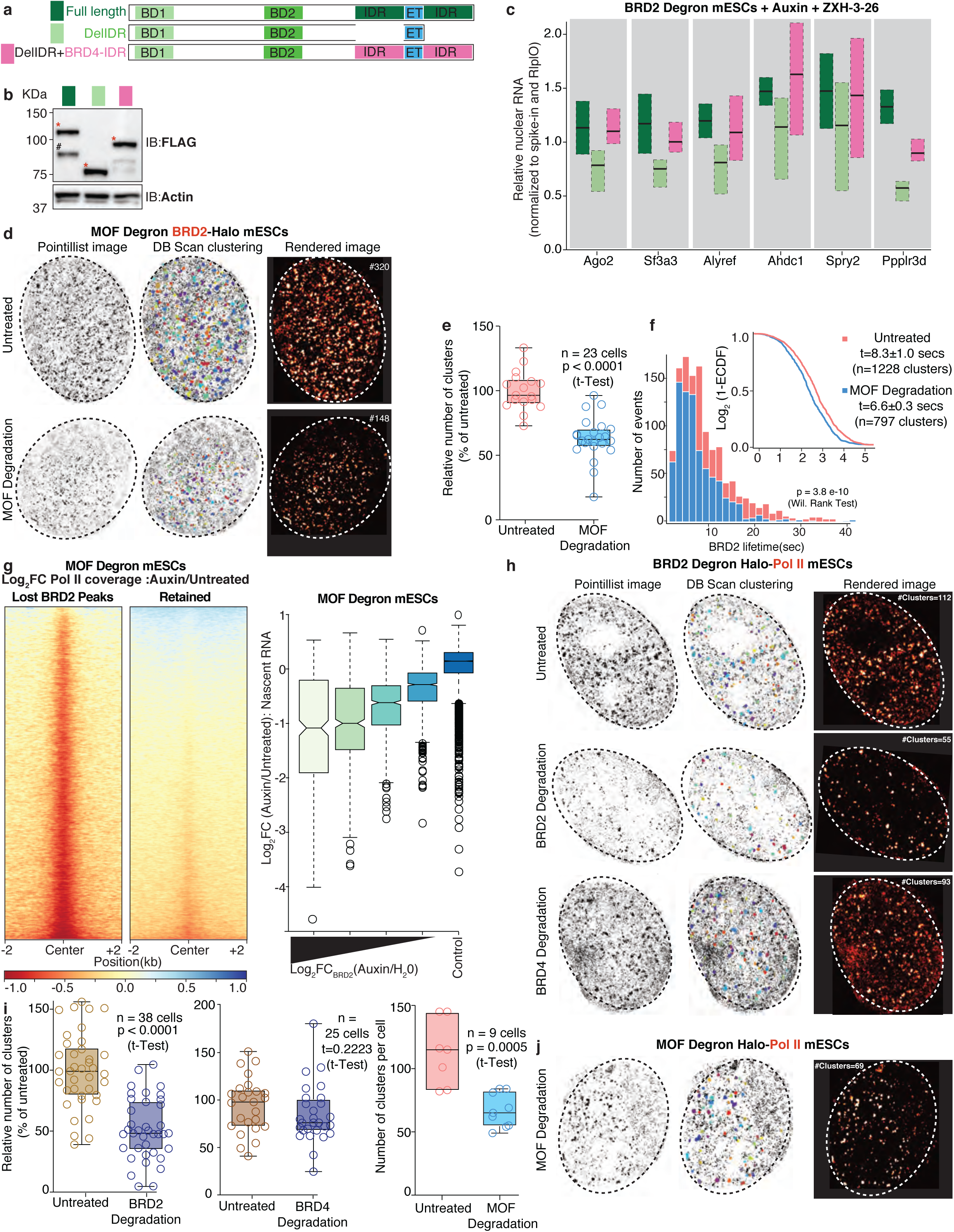
MOF-mediated H4 acetylation is crucial for the spatial organization and function of BRD2 in regulating transcription dynamics. **a**, Schematic depicting the domain organization of BRD2 variants. All the mutants carry the two BRD2 bromodomains and the BRD2 ET domain. They only differ in their IDRs. For replacement of the BRD2 IDR with an exogenous IDR, a segment of the BRD4 IDR with a comparable size to the original BRD2 IDR was used. **b**, Western blot showing the expression of 3xFLAG-tagged BRD2 variants in cells. Red asterisks denote the specific bands. Actin serves as a loading control. **c**, Boxplots depicting the changes in nuclear RNA levels of the mentioned transcripts upon expression of BRD2 variants in the absence of endogenous BRD2 and BRD4. **d**, Example of super-resolved live-cell images of BRD2 in BRD2-Halo MOF degron mESCs before and after auxin treatment. **e**, Boxplot depicting the changes in the number of BRD2 clusters upon MOF depletion as a percentage of the untreated condition (*n*=23 cells). **f**, Histogram (left) and empirical cumulative distribution function (ECDF) plot (right) showing the distribution of BRD2 cluster lifetime before and after rapid MOF depletion. **g**, ChIP-Seq analysis depicting the log2FC of RNA Pol II coverage upon 3 h of auxin treatment of MOF degron mESCs for 3 h. The signal was plotted over the BRD2 ChIP-Seq peaks that were lost or retained upon MOF depletion **(left)** and boxplot depicting the log2FC of nascent RNA levels in relation to differential BRD2 enrichment over their promoters upon rapid MOF depletion. All expressed genes which exhibited differential BRD2 binding at their promoters following MOF depletion were sorted into 4 quartiles of equal sizes based on the log2FC of BRD2 occupancy upon MOF depletion. The expressed genes with no differential BRD2 enrichment at their promoters served as a control (dark blue) **(right)**. **h, j**, Example of super-resolved live-cell images of Halo-RPB1 in the corresponding degron mESCs with endogenous Halo-RPB1 upon BRD2, BRD4 **(h)** or MOF depletions **(j)**. **i**, Boxplot depicting the changes in the number of RNA Pol II clusters upon BRD2, BRD4 and MOF depletion as percentage of untreated condition.

Given the fact that the formation of BRD2 foci upon inhibition of pause release depends on BRD2’s chromatin binding, we next asked whether H4 acetylation might regulate the dynamics of BRD2 transient clusters. To this end, we have endogenously tagged BRD2 with a Halo tag in MOF degron mESCs using CRISPR-Cas9 and performed live-cell super-resolution microscopy. Rapid depletion of MOF resulted in a ca. 50 % decrease in the number of BRD2 transient clusters and their overall destabilization as evident by their reduced average half-life (Untreated: 8.3 ± 1 s, MOF-depleted: 6.6 ± 0.3 s) (Fig. 5d-f). Along with our previous BRD2 ChIP-Seq results upon MOF depletion (Fig. 1e), this finding implies that histone acetylation does not only control the chromatin binding, but also the spatial organization of BRD2.

We were interested in understanding the functional consequences of the loss of BRD2 chromatin binding and subsequent failure of the assembly of BRD2 transient clusters. We therefore performed TT-Seq and RNA Pol II ChIP-Seq experiments in MOF degron mESCs to analyze the transcriptional dysregulation resulting from MOF depletion in relation to MOF-sensitive BRD2 peaks. Rapid MOF depletion resulted in significant transcriptional downregulation of 1,086 genes and its target genes exhibited reduced RNA Pol II levels at their promoters (Extended Data Fig. 2j, k). In line with our previous observations (Fig. 3a-c), MOF depletion led to more severe reductions in RNA Pol II coverage around MOF-sensitive BRD2 peaks in comparison to that around MOF-insensitive BRD2 peaks (Fig. 5g, left). Furthermore, BRD2 loss upon MOF depletion correlated with a more severe reduction in RNA synthesis (Fig. 5g, right).

Our results so far collectively imply that BRD2 depletion or loss of its chromatin binding alters the enrichment of RNA Pol II at the promoters in a rapid manner. Since genomics approaches characterize RNA Pol II at a population level and steady state without fully capturing the dynamics of the transcription machinery, we performed live-cell super-resolution microscopy experiments to study the real-time responses of RNA Pol II to rapid BRD2 depletion. To this end, we have endogenously tagged the largest subunit of RNA Pol II, Rpb1, at its N-terminus with a Halo tag using CRISPR-Cas9 in BRD2 and MOF degron mESCs and analyzed the dynamics of sub-diffraction RNA Pol II clusters, which are known to be associated with transcription initiation ^21^. Similar to the previous reports, tcPALM experiments identified RNA Pol II clusters with a median lifetime of 6.5-7.5 seconds (Fig. 5h and Extended Data Fig. 3g) ^21^. Depletion of BRD2 alone upon auxin treatment for 3 h led to an average reduction of ca. 50 % of all RNA Pol II clusters. Interestingly, the half-life of the remaining RNA Pol II clusters upon BRD2 depletion did not show any significant differences, possibly indicating that not all RNA Pol II clusters depend on BRD2 (Fig. 5h, i and Extended Data Fig. 3g). Surprisingly, short-term depletion of BRD4 did not significantly affect RNA Pol II cluster formation, but rather increased the half-life of existing clusters, further emphasizing the roles of BRD2 and BRD4 in related, but different layers of transcription regulation (Fig. 5h, i and Extended Data Fig. 3g). Similarly to BRD2 depletion, acute loss of MOF also reduced the number of RNA Pol II clusters, providing further evidence that MOF and BRD2 work on the same axis to regulate transcription initiation dynamics (Fig. 5h-j and Extended Data Fig. 3g).

## Discussion

Within the BET BRD protein family, the function of BRD4 is best understood given its interaction with pTEFb complex, the strong reduction in transcription elongation upon its degradation and its association with oncogene expression ^18,22,23^. In contrast, other widely expressed BET BRD members, BRD2 and BRD3 have often been neglected since their depletion alone does not lead to obvious defects in global transcription ^4,18^. In this study, we have systematically dissected the function of BRD2 as a reader of histone H4 acetylation and its hierarchical interplay with other BET BRD members. Our unique approach integrating population-based genomics and single-cell imaging analyses unraveled a previously unknown role of BRD2 in maintaining chromatin occupancy of RNA Pol II at the promoters, which becomes more crucial in the absence of BRD4 or upon inhibition of pause release (Extended Data Fig. 3h).

By performing co-depletion of BET BRD proteins combined with nascent RNA and RNA Pol II ChIP-seq experiments as well as super resolution microscopy, we demonstrate a clear functional segregation of BET BRD proteins in transcription regulation: While BRD4 acts as an essential regulator of transcription elongation by promoting the release of RNA Pol II from promoter-proximal pausing, BRD2 and potentially BRD3 protect the cell from a total collapse of global RNA synthesis in the absence of BRD4. This unique view of BET BRD members as distinct, but collaborative functional layers, also provides an explanation why BRD2, BRD3 and BRD4 usually co-exist at the same chromatin binding sites.

Despite the overlap in the genome-wide chromatin binding profiles of BRD2, BRD3 and BRD4, it was surprising that only BRD2 predominantly responds to the loss of H4-ac mediated by MOF. This was further corroborated by a recent proteomic profiling study of chromatin readers suggesting that BET BRD proteins preferentially bind to H4-acetylated rather than H3-acetylated nucleosomes ^6^. Considering the dramatic loss of BRD2 chromatin binding upon MOF depletion, it is very likely that the specificity of BRD2’s bromodomain modules may more strongly depend on H4K16ac and its combination with other H4-ac marks in comparison to BRD3 and BRD4 ^5,28^. Since these proteins almost exclusively associate with actively transcribed regions which are enriched for both H4-ac and H3-ac, their combination might be potentially maintaining the chromatin association of BRD3 and BRD4 due to an avidity effect. Accordingly, their loss of chromatin binding might have required the single or combined depletion of other HATs.

Our integrative approach combining genomics and super-resolution microscopy revealed that H4-ac is not only essential for BRD2 recruitment to chromatin, but also dictates its spatial organization in the nucleus. BRD2 is organized into transient clusters under steady-state conditions while it forms dynamic foci upon inhibition of RNA Pol II pause-release. We demonstrated H4 acetylation as a prerequisite for BRD2 function by depleting the histone acetyltransferase MOF. Furthermore, we observed increased chromatin binding of BRD2 upon inhibition of pause-release. These differential chromatin binding properties of BRD2 provide compelling evidence that chromatin binding of BRD2 is essential for its condensation dynamics. Thus, our findings collectively provide examples of how the chromatin landscape and the transcriptional state can target transcriptional condensates to specific loci.

Mechanistically, our data suggest that there is a dynamic interplay between promoter-proximal RNA Pol II, BRD2 and BRD4. In the presence of BRD4 and very likely other transcriptional coactivators promoting rapid pause release, the function of BRD2 in promoting transcription initiation and thereby chromatin association of paused RNA Pol II becomes less pronounced since a rapid transition into productive elongation takes place. By depleting BRD4 or inhibiting pause release using flavopiridol, we managed to take snapshots of the dynamic relationship between RNA Pol II and BRD2 and could uncover the important role of BRD2 in the control of RNA Pol II dynamics at the promoters. The accumulation of BRD2 on chromatin upon pause release inhibition and the appearance of dynamic foci suggests that the clustering dynamics of BRD2 might be mechanistically relevant. In particular, the correlation between the severity of RNA Pol II loss from promoters upon BRD2 depletion and the increase in BRD2 chromatin occupancy upon pause release inhibition supports this idea. Furthermore, we also provide direct evidence that BRD2 clustering is required for its function by using BRD2 variants differing in their IDR. Nevertheless, how inhibition of pause release promotes chromatin localisation or retention of BRD2 remains to be elucidated. However, the previously reported increase in BRD2 chromatin occupancy upon BRD4 depletion further supports the notion that BRD2 acts as a sensor of prolonged RNA Pol II pausing ^24^.

Even though we majorly focused on the functional relationship between BET proteins in transcription regulation, they also have been proposed to have distinct roles in the regulation of genome architecture: It has been recently demonstrated that BRD2 contributes to the compartmentalization of active chromatin upon cohesin loss in an antagonistic manner with BRD4 ^24^. Nevertheless, the impact of BRD2 depletion alone on genome compartmentalization was less prominent ^24,25^. Interestingly, BRD4 has also been shown to orchestrate genome folding during neural crest differentiation by its interaction with cohesin cofactor NIPBL ^26^. In this respect, the dynamic interplay between BET proteins and cohesin in genome folding throughout physiological and potentially pathological cellular states remains to be further studied.

Collectively, our study focusing on the BET proteins provides an important example for the functional role of acetylation-dependent transcriptional coactivator condensation in transcription dynamics. Our data supports a model by which the differential condensation properties of BRD2 and BRD4 give rise to distinct modules which support transcription initiation and elongation, respectively. The synergistic roles of BRD2 and BRD4 in transcription control also imply a crosstalk on the level of their condensation dynamics. Given the prominence of BET inhibitors in preclinical studies, our work provides a new mechanistic avenue to consider to further develop their therapeutic efficacy.

## Acknowledgments

We thank all MPI-IE facilities for technical support, particularly imaging, FACS, deep-sequencing, mass spectrometry and bioinformatics facilities. We acknowledge all the past and present members of Akhtar lab for providing critical feedback. We are grateful to M. Wiese and Y. Zhou for critical reading of the manuscript. We thank M. Du and C. Lee for sharing reagents and providing feedback related to super-resolution microscopy.

## Author Contributions

N.U.E. and A.A. conceptualized the study. N.U.E. designed the experiments and performed transcriptomics, genomics, CRISPR-Cas9 genome editing, biochemistry experiments and analyzed all the data. S.G. performed live-cell confocal and super-resolution imaging and analyzed all imaging data with the help of J.A.M.L. and E.M.P. R.O. helped with TT-Seq data analysis and provided critical feedback on TT-Seq experiments. I.I.C. supervised E.P. and provided critical feedback on super-resolution microscopy experiments. A.A. supervised N.U.E., S.G., R.O. and acquired funding. N.U.E., S.G and M.S. wrote the manuscript with input from all the authors.

## MATERIALS AND METHODS

### Cell culture

WT26 male mouse embryonic stem cells (ESCs) were a kind gift of Thomas Jenuwein. ESCs were cultured at 37°C and 5 % CO2 on 0.1% gelatin (Merck #G1393)-coated dishes and were maintained in Dulbecco’s Modified Eagle’s Medium (DMEM, Gibco #37966-021) supplemented with 15% heat-inactivated ES-cell specific fetal calf serum (FCS), 100 U/mL penicillin and 100 U/mL streptomycin (Gibco #15140-122), 1mM glutamine (GlutaMax, Gibco #35050061), 1mM sodium pyruvate (Gibco #11360-70), non-essential amino acids (NEAA, Gibco #11140050), ß-Mercaptoethanol (Gibco #31350-010), and in-house purified LIF. 90% FCS and 10% DMSO (Sigma #8418) was used for freezing the ESCs for storage. All the cell lines were regularly tested negative for mycoplasma contamination.

### Generation of Tir1-expressing stable cell line

For stable expression of Tir1, the reporter TALEN system was used as previously described ^27^. Briefly, a genome cassette containing a CAG promoter, cDNA sequence of *Tir1* along with 3xMyc-HA tag and a BGH-poly(A) signal, was cloned into a targeting vector (a kind gift of Bühler lab) which contains the homology arms for insertion into *Rosa26* locus. This repair template along with pRR-Puro reporter and heterodimeric ELD/KKR *FokI* domain plasmids targeting *Rosa26* locus were co-transfected into WT mouse ES cells using Lipofectamine 2000 (Invitrogen #11668027) according to the manufacturer’s instructions. One day after transfection, ESCs were selected using 2 μg/mL puromycin (Gibco #A11138-03) for 36 hours. Positive clones were picked using pipette tips and transferred into 96-well plates for further screening of Tir1 protein level.

### CRISPR-Cas9 genome editing

In order to endogenously tag proteins of interest with tags at their N- or C-termini, CRISPR-Cas9 genome editing was used. Guide RNAs were designed using CRISPR web toolbox CHOPCHOP ^28^.

For phosphorylation and annealing, 1 μL each of 100 μM forward and reverse oligonucleotides containing the guide RNA sequences with the appropriate extensions for cloning were mixed with 1 μL of T4 PNK Kinase (NEB #M0201), 1 μL of 10 mM ATP and 1 μL of 10X T4 DNA Ligase buffer in total volume of 10 μL. This reaction was first incubated at 37°C for 30 minutes, then at 95°C for 5 minutes and finally cooled down to room temperature with a ramping rate of 0.1°C/s. The annealed oligos were diluted 1/200 in ddH2O and 2 μL of this dilution was used for ligation into 100 ng of PX458 or PX459 backbones, that have been digested with *BbsI* (NEB #r3539) and gel-purified, using T4 DNA Ligase (NEB #M0202).

In order to generate homology-directed repair (HDR) templates, 5’- and 3’ homology arms were designed adjacent to the cut site of Cas9 with a minimum length of 750 bp. Inserts for protein tagging were obtained either from gene synthesis or from plasmids that were previously generated in the lab. The homology arms, PCR-amplified from mouse genomic DNA, and the inserts were assembled together into pJET1.2 backbone using Gibson Assembly (NEB). HDR template and sgRNA for N-terminal endogenous tagging of RPB1 with Halo were a kind gift of Cisse lab ^29^.

3 μg of HDR template and 1 μg of sgRNA-containing Cas9 backbone were co-transfected into 1x106 ESCs (seeded in each well of a 6-well plate) using lipofectamine 2000 according to the manufacturer’s protocol. 6 hours after the transfection, the culture medium was exchanged and the ESCs were allowed to recover for at least another 6 hours under standard growth conditions. Then the ESCs were re-plated on a 10 cm plate and selected with appropriate antibiotics for 4-7 days. The positive clones were picked using pipette tips and transferred into 96-well plates for PCR screening and further growth. For proteins that were endogenously tagged with a fluorescent marker, the above described strategy was used with the difference that single ESCs positive for the fluorescent tag were FACS sorted into 96-well plates.

### Small molecule treatments

To induce rapid degradation in the degron cells, they were treated with 500 μM of auxin (3-indole acetic acid, Sigma) in ddH2O. We have used the following inhibitor concentrations for the rest of the compounds: Flavopiridol (10 μM in DMSO, Sigma), ZXH-3-26 (50 nM in DMSO, Tocris Bioscience), dBET6 (100 nM, Tocris Bioscience), DRB (100 μM in DMSO, Sigma), JQ1 (1μM in DMSO, Sigma), Actinomycin D (25 μg/mL in DMSO, Sigma).

### Immunoblotting

Cellular pellet was resuspended and lysed in SDS loading dye (2X ROTI-Load , Carl Roth). Samples were boiled at 95°C for 5 minutes, sonicated using the Branson sonifier 250 (40% duty cycle, 1.5 output, 10 pulses) and boiled again for 5 minutes. SDS-PAGE was performed by running samples on NuPAGE Bis-Tris 4-12 % gradient gels (Invitrogen #NP0321PK2) using MOPS (Novex #NP0001, for detection of proteins with molecular weight higher than 50 KDa) or MES buffer (for detection of proteins with molecular weight lower than 50 KDa, for example histones).

For detection of proteins with molecular weight lower than 30KDa (for example, the histones), proteins were wet-transferred to a 0.22 uM PVDF membrane, otherwise a 0.45 uM membrane was used. Following the transfer, the membrane was blocked with 5% skimmed milk in 0.3% Tween-PBS for 1 hour at RT and incubated with relevant primary antibodies overnight. After washing thrice for 5 minutes each with 0.3% Tween-PBS, membranes were incubated with HRP-conjugated secondary antibodies for 1.5 hours at RT. The membrane was washed thrice for 5 minutes each with 0.3% Tween-PBS and finally developed with Lumi light enhanced chemiluminescence substrate (Roche #12015196001) and imaged using BioRad ChemiDoc system.

### Protein purification and *in vitro* assays

Design of the construct, purification process of the protein and *in vitro* droplet assay was performed according to (Sabari *et al.* 2018) with minor modifications ^20^. BRD2-IDR was amplified from cDNA and the eGFP tag was incorporated at its 5’ end. The recombinant BRD2-IDR-eGFP was cloned into pET41 expression vector and transformed into BL21(DE3)LysS bacterial strain. Cells were grown in LB containing kanamycin and chloramphenicol at 37°C till the OD of 0.6-0.7 and then 1 mM IPTG was added to allow protein expression at 18°C overnight. The bacterial culture was collected by centrifugation and the pellet was resuspended in Buffer A (50mM Tris pH7.5, 500 mM NaCl) containing 10mM imidazole, cOmplete protease inhibitors (Roche, 11873580001) and sonicated (10 cycles of 15 seconds on, 60

seconds off). The lysate was clarified by centrifugation at 12,000g for 30 minutes at 4°C and added to Ni-NTA agarose slurry (Invitrogen, R901-15), pre-equilibrated with 10X volumes of Buffer A. The recombinant protein was allowed to bind to the column through its His-tag at 4°C for 1.5 hours, with gentle rotation. The agarose slurry was poured into a column, packed, and washed with 15x volumes of Buffer A containing 10mM imidazole. Protein was eluted in multiple fractions with Buffer A with 250mM imidazole. Fractions of elutions containing protein as judged by coomassie stained gel were combined and dialyzed against Buffer B (50mM Tris-HCl pH7.5, 500 mM NaCl, 10% glycerol, 1mM DTT).

Recombinant BRD2-IDR-eGFP was concentrated and desalted to an appropriate protein and salt concentration using Amicon Ultra centrifugal filters (30K MWCO, Millipore). Solutions containing varying concentrations of the recombinant protein, salt, RNA or ATP were prepared, incubated for 30 minutes at RT and loaded onto glass slides and mounted with cover slips. Slides were then imaged using an inverted confocal laser scanning microscope Zeiss LSM 880 equipped with an Airyscan detector.

### BRD2 interactome

In order to minimize the saturation of MS peaks with antibody peptides used for co-immunoprecipation, antibodies were crosslinked to magnetic DynaBeads using DMP: Per IP, 50 µL of resuspend Dynabeads was washed twice with 500 µL of 0.1 M phosphate buffer (pH 8.0) by inverting and gentle vortexing. Then, the beads were first incubated with 6 µg of the antibody in phosphate buffer (pH 8.0) in a total volume of 100 µL for 30 minutes at 4 °C. Afterwards, Tween-20 was added to 0.1 % and the sample was further incubated at 4 °C overnight. The next day, the beads were first washed thrice in 500 µL of phosphate buffer and then twice in 500 µL of 0.2 M triethanolamine. The antibody was crosslinked to the beads by incubation with 500 µL DMP (6.5 mg/mL in 0.2 M triethanolamine) for 45 minutes at room temperature. The crosslinked beads were rinsed once with 500 µL of 0.1 M ethanolamine and then incubated in 0.1 M ethanolamine for 30 minutes at room temperature. After washing thrice with 500 µL of PBS, the excess non-crosslinked antibody was washed away by incubating the beads in 500 µL of 0.1 M Glycine-HCl (pH 2.5) by inverting and gentle vortexing. The beads were then washed twice with 1X PBS and finally resuspended in 100 µL of 1X PBS supplemented with 0.1 % Tween-20 and 0.02 % Na-Azide. The beads were stored at 4 °C until use.

1 x 10^7^ cells were harvested per replicate using accutase and flash-frozen. The cell pellet was thawn on ice and lysed in 2 mL of Buffer A (10 mM HEPES-KOH, pH 7.9, 5 mM MgCl_2_, 10 mM KCl, 1 mM DTT, 0.1 % NP-40) by rotating for 10 minutes at 4 °C. The nuclei were collected by spinning down the sample for 10 min at 2000 rpm at 4 °C. The nuclear pellet was resuspended in 100 µL of buffer B (25 mM HEPES-KOH, pH 7.5, 150 mM KCl, 10 % Glycerol (v/v), 12.5 mM MgCl_2_, 0.2 % NP-40, 5 mM CaCl_2_) and kept on ice for 10 minutes. Then, 2 µL of MNase was added and the sample was incubated for 10 min at 37 °C in a thermomixer at 1000 rpm. MNase digestion was quenched by addition of 0.73 µL of 330 mM EGTA. The soluble nuclear fraction was collected by spinning down the samples at 16000 g for 10 min at 4 °C. The volume of the samples brought to 500 µL using IP buffer (25 mM HEPES-KOH, pH 7.5, 150 mM KCl, 10 % Glycerol (v/v), 12.5 mM MgCl_2_, 0.2 % NP-40, 31.25 mM EDTA, 31.25 mM EGTA). The crosslinked beads were washed thrice in IP buffer and then incubated with the samples overnight at 4 °C. The next day, beads were washed three times with 500 µL of IP buffer and three times with 500 µL of WB2 buffer (25 mM HEPES-KOH, pH 7.5, 150 mM KCl, 5 % Glycerol (v/v), 5 mM MgCl_2_, 0.02 % Rapigest-MS).

For on-bead digestion of the samples for LC-MS/MS, the beads were resuspended in 50 µL of 2TU/0.04% Rapigest-MS and TCEP was added to 10 mM. After incubation for 5 minutes at 25 °C, CAA was added to 40 mM. After incubation for another 5 minutes, IPed proteins were digested with 0.2 µg trypsin and 0.2 µg LysC for 90 minutes at 25 °C. Afterwards, digested peptides were transferred into a fresh tube. Magnetic beads were further incubated with 50 mM ABC for 10 minutes at 25 °C and the left-over peptides were also collected. The pool of peptides were further digested with 0.28 µg of trypsin at 25 °C overnight. After treating with 10 µL of 10 % TFA, the samples were mixed with 50 µL of stage A buffer (0.5 % AcOH, 0.02 % HFBA) . After spinning down for 10 min at 14000g, the samples were loaded onto C18 STAGE tips for further processing for LC-MS/MS.

### ChIP-Seq

ESCs were crosslinked with 1 % Formaldehyde in 1X PBS for 10 minutes at RT followed by quenching with 125 mM Glycine at 4 °C. After washing the ESCs twice with PBS, ESCs were lysed in LB1 buffer (50 mM HEPES, pH 7.9, 140 mM NaCl, 1 mM EDTA, 10 % Glycerol, 0.5 % NP-40, 0.25 % Triton X-100) for 20 minutes at 4 °C. Then, samples were washed in LB2 buffer (10 mM Tris-HCl, pH 8.0, 200 mM NaCl, 1 mM EDTA, 0.5 mM EGTA) for 5 minutes at 4 °C. Chromatin was extracted by incubation in LB3 buffer (10 mM Tris-HCl, pH 8.0, 100 mM NaCl, 1 mM EDTA, 0.5 mM EGTA, 0.1 % DoC-Na, 0.5 % Na-Sarcosinate, 1 % Triton X-100) for 10 minutes on ice followed by sonication for 12 minutes on a Covaris sonicator. The sonicated chromatin was spun down at 16,000g for 10 minutes at 4 °C and the supernatant was incubated with the appropriate antibody overnight at 4 °C. Antibody-bound chromatin fragments were collected by incubation with Protein A or G DynaBeads for 3 h at 4 °C. The beads were washed for 5 minutes at 4 °C with the following buffers: RIPA-150 (50 mM Tris-HCl, pH 8.0, 150 mM NaCl, 0.1 % SDS, 0.5 % DoC-Na, 1 % NP-40), RIPA-500 (50 mM Tris-HCl, pH 8.0, 500 mM NaCl, 0.1 % SDS, 1 % NP-40), Li-Buffer (50 mM Tris-HCl, pH 8.0, 250 mM LiCl, 0.5 % DoC-Na, 1 % NP-40), TE Buffer (50 mM Tris-HCl, pH 8.0, 10 mM EDTA). After an additional wash with TE Buffer, beads were resuspended in 100 μL of elution buffer (50 mM Tris-HCl, pH 7.4, 1 % SDS, 1 mM EDTA) and crosslinks were reversed by incubation at 65 °C for at least 4 h. Proteins and RNA were degraded by further incubations with RNase A and Proteinase K, respectively. Resulting DNA was purified using ChIP DNA Clean and Concentrator kit (Zymo) according to the manufacturer’s protocol. 1-10 ng of purified Input and ChIP DNA was used to prepare Illumina multiplexed sequencing libraries using NEBNext Ultra II DNA Library Prep Kit.

### Native histone ChIP-Seq

3 x 10^6^ ESCs per replicate were harvested using accutase, washed once in 1X PBS and flash-frozen. The cell pellet was thawn on ice and resuspended in 150 μL of 1X PBS and mixed with 150 μL of 2X Lysis buffer (100 mM Tris-HCl, pH 8.0, 0.2 % Triton X-100, 0.1 % DOC-Na, 10 mM CaCl2, 10 mM Na-Butyrate) containing 0.9 μL of MNase. The cell lysate was incubated first for 20 minutes at 4 °C and then for 10 minutes at 37 °C. The MNase digestion was quenched by addition of 30 μL of 25 mM EGTA. Afterwards, the sample was immediately placed on ice and mixed ocassionally by inverting the tube for 2 minutes. The soluble chromatin was collected by spinning down the sample at 16000g for 5 minutes at 4 °C and mixed with the dilution buffer (50 mM Tris-HCl, pH 8.0, 150 mM NaCl, 0.1% DOC-Na, 2.2% Triton X-100, 50 mM EDTA, 50 mM EGTA, 5 mM Na-Butyrate). 2 % of the soluble fraction was taken as input and the rest was incubated overnight with the histone PTM antibody at 4 °C. The next day, magnetic Protein A or G Dynabeads were washed in PBS thrice and 20 μl per sample were added to collect the antibody-chromatin conjugates. Then, they were washed in the following buffers, each for 5 minutes at 4 °C: Low salt buffer (0.1 % SDS, 1 % Triton X-100, 0.1 % DOC-Na, 20 mM Tris-HCl, pH 8.0, 2 mM EDTA, 150 mM NaCl)[twice], high salt buffer (0.1 % SDS, 1 % Triton X-100, 0.1 % DOC-Na, 20 mM Tris-HCl, pH 8.0, 2 mM EDTA, 360 mM NaCl), LiCl buffer (0.25 M LiCl, 1 % NP-40, 1 % DOC-Na. 1 mM EDTA, 10 mM Tris-HCl, pH 8.0) [twice]. Finally, the samples were rinsed once in TE buffer (50 mM Tris-HCl, pH 8.0, 10 mM EDTA) and resuspended in 100 μL of elution buffer (50 mM Tris-HCl, pH 7.4, 1 % SDS, 1 mM EDTA) and incubated first for 1 h at 55 °C with Proteinase K and then for 30 minutes at 37 °C with RNAse A. Resulting DNA was purified using ChIP DNA Clean and Concentrator kit according to manufacturer’s protocol. 1-10 ng of purified Input and ChIP DNA was used to prepare Illumina multiplexed sequencing libraries using NEBNext Ultra II DNA Library Prep Kit.

### Total RNA-Seq

For total RNA-Seq and Quantitative RT-PCR (RT-qPCR) experiments, ESCs seeded on 6-well plates were directly lysed on plate using 300 μl of Trizol reagent (Invitrogen). This lysate was either stored at -80 °C or immediately processed for RNA purification using Direct-Zol RNA Miniprep Kit (Zymo Research) according to the manufacturer’s instructions. RNA concentration was measured using Qubit 2.0. Total RNA of high quality was used for library preparation using Illumina Stranded TruSeq Total RNA RiboZero Plus kit.

### TT-Seq

TT-Seq was performed as previously described with minor modifications ^17^: ESCs were treated with 500 μM 4-sU for 5 minutes under regular growth conditions and lysed using Trizol (Invitrogen) immediately afterwards. Total RNA was purified using chloroform and isopropanol precipitation and resuspended in TE buffer. 300 μg of total RNA for each sample was spiked with spike-in mix (appendix) and fragmented using Bioruptor Plus (30 seconds ON, 30 seconds OFF, 1 cycle, HIGH setting) and an aliquot was taken as Total RNA sample. The rest of the fragmented total RNA was biotinylated using Biotin-HPDP (Thermo Fisher) at room temperature for 2 hours. To remove excess Biotin-HPDP, RNA was precipitated using phenol:chloroform-isoamyl alcohol (Sigma) and ethanol and resuspended in ddH2O. Biotinylated RNA fragments were purified using μMACS streptavidin beads and μMACS columns (Miltenyi Biotec) according to the recommendations of the manufacturer. Labeled RNA was eluted twice using 100 mM DTT and purified using Oligo Clean and Concentrator Kit (Zymo Research). Both total and labeled RNA samples were treated with Turbo DNAse (Invitrogen) for 30 minutes at 37 °C and purified again Oligo Clean and Concentrator Kit (Zymo Research). Libraries were prepared using Illumina Stranded Total RNA with RiboZero Plus according to the manufacturer’s instructions.

### Confocal microscopy

For immunofluorescence, ESCs grown on coverslips to a confluency of 70-80% were fixed with 4% formaldehyde, diluted in PBS, at RT for 15 minutes and permeabilized with 0.2% Triton X-100-PBS for 10 minutes. The ESCs were then blocked with 4 % BSA, prepared in 0.05% Triton X-100-PBS for 1 hour and then incubated overnight with primary antibodies in the above blocking buffer at 4°C. Next day, the ESCs were washed thrice at RT, 10 mins each, with 0.05% Triton X-100-PBS and then incubated with fluorophore-conjugated secondary antibodies for 2 hours at RT in the dark.Then they were washed twice, 10 mins each, with 0.05% Triton X-100-PBS and then counterstained with 20 μM Hoechst 33342 in PBS for 10 minutes. The coverslips were mounted on glass slides with using FluoroGel (#GTX2814) after a final PBS wash of 10 minutes.

For steady and time lapse live cell imaging, ESCs were grown on eight-well glass bottom Ibidi chambers 6-8 hours before the start of the experiment. Experiments were performed using inverted confocal laser scanning microscope Zeiss LSM 880 equipped with an Airyscan detector. The laser power was set to a maximum of 1% to minimize photobleaching and phototoxicity. Images were acquired at different focal planes with appropriate Z-stack settings to cover the width of the nucleus.

Fluorescence Recovery After Photobleaching (FRAP) experiments were performed as follows. 1-3 equal sized regions of interest in a cell were selectively bleached with 100% laser power and mages were collected each 2 seconds. Fluorescence intensity of the regions of interest was calculated and corrected against the mean of 2 same sized regions containing unbleached clusters for the corresponding field of view. Image acquisition and analysis were performed using Zen2 Blue Software (version 3.1 and 3.2) by Zeiss.

### Super-resolution imaging

Live-cell Photoactivated Localization Microscopy (PALM) was carried out on ESCs expressing Halo-tagged protein of interest. 100,000 ESCs were seeded the night before the experiment on 35 mm MatTek imaging dishes which were pre-coated with 5 ug/ml poly-L-Ornithine (PLO, Sigma) (for at least 5 hours) and 10 ug/ml mouse Laminin (for at least another 5 hours). On the day of the experiment, ESCs were incubated with 100nM photoactivatable halo ligand, PA-JF646, for 10 minutes, washed twice with imaging media followed by a further 20 minutes incubation to wash out the unbound ligand. Cells were simultaneously illuminated with low intensity near UV light (405 nm), which was gradually increased from 0.1% to 10% over the time of the acquisition, for photoactivation of Halo ligand for fluorescence detection with an exposure time of 50 ms and images were acquired for 10,000 frames.

Image analysis was performed using the qSR tool^30^. Density-Based Spatial Clustering of Applications with Noise (DBSCAN) embedded in the qSR software was used to define clusters with a minimum 25 nearby localizations within 100 nm. Transient clusters were manually marked and verified for calculating the cluster lifetime.

### ChIP-Seq analysis

ChIP-Seq data was processed using the default parameters of SnakePipes v. 2.5.1 DNA-mapping and ChIP-Seq pipelines with “--trim --dedup --mapq 3 --properPairs” options ^31^. For data visualization, replicates with a Pearson correlation R>0.9 are merged. Heatmaps and metagene plots have been generated using deeptools ^31,32^. Custom R scripts have been used for the calculation of ChIP-Seq RPKM over different regions. To perform differential binding analysis, R Diffbind package was used ^33^. The genome snapshots were generated using pyGenomeTracks ^34^.

### Total RNA-Seq analysis

Total RNA-Seq was processed using default parameters of SnakePipes v.2.5.1 noncoding-RNA-seq pipeline with “--trim” option ^31^. Downstream analysis and visualization was performed using featureCounts (v.2.0.0), and R packages DESeq2 (v.1.34.0), ggplot2 (v.3.3.5) ^35–37^. Volcano plots were produced using EnhancedVolcano package in R (<u>github.com/kevinblighe/EnhancedVolcano</u>). For time-course experiments, unless stated otherwise, the ’untreated’ condition was always taken as reference.

### TT-Seq analysis

TT-Seq data analysis was performed as previously described ^38^. The data were processed through the snakePipes mRNA-seq pipeline (modified version of v.2.1.2), similar to RNA-seq analysis. Adapters and low-quality bases (<Q20) were removed using Cutadapt (v.2.8; https://github.com/marcelm/cutadapt). For all of the samples, reads were then mapped to the reference genome with STAR (v.2.7.4a) ^39^. Then, reads per gene were counted using featureCounts (v.2.0.0) ^36^. The gene-level counts obtained from featureCounts were then used for differential expression analysis using DESeq2 (v.1.26.0), similar to the RNA-seq analysis ^35^. For transcriptional rate calculations, read counts for all genes were obtained from each corresponding labeled and unlabelled TT-seq sample. To estimate the rates of RNA degradation and synthesis, we used a statistical model that was described previously ^17^.

## Data availability

All the NGS datasets and differential expression analysis tables generated in this study will be deposited to the Gene Expression Omnibus.

## Code availability

We have made use of publicly available softwares, tools and R scripts for data analysis. Specific codes used for the analysis of NGS datasets in this manuscript are available upon request.

