## Supplementary Figures for "Acetylation-dependent clustering of BRD2 instructs transcription dynamics"

Extended Data Fig. 1

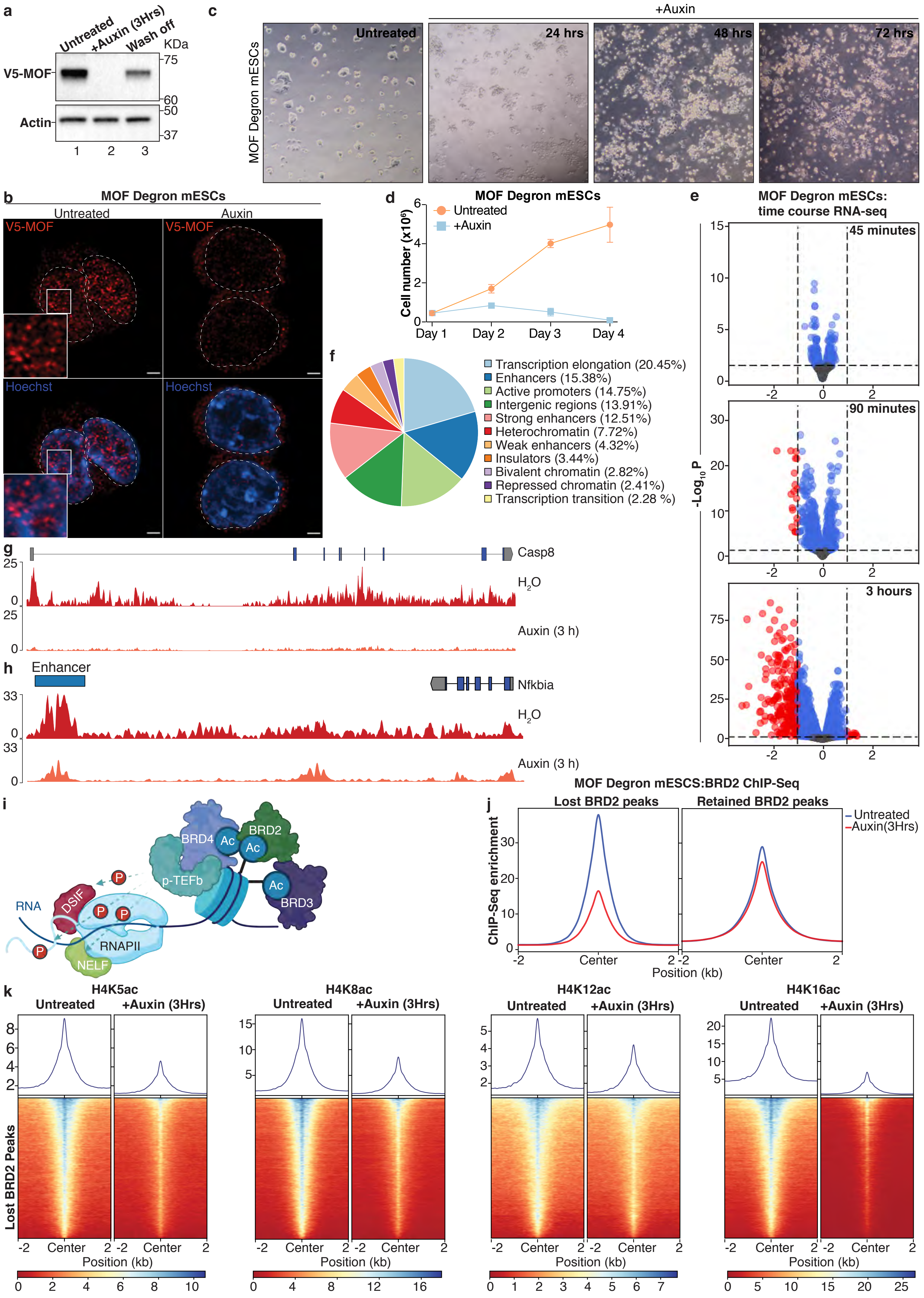

**Extended Data Fig. 1: Characterization of MOF degron mESCs and BRD2 chromatin binding in association with H4-ac.**

**a,** Immunoblot analysis of MOF degron mESCs left untreated, treated with auxin for 3 hours ("+Auxin (3Hrs)") or treated with auxin for 3 hours followed by wash-off of auxin and incubation in auxin-free media for 12 hours ("Wash off"). **b,** Immunofluorescence microscopy images of untreated and auxin-treated (12 hours) MOF degron mESCs using an antibody against the V5-tag (red) and nuclear Hoechst staining (blue). Dashed line represents the nuclear boundary. Scale bar of images: 5 μm. **c,** Phase contrast microscopy images of untreated and auxin-treated MOF degron mESCs to qualitatively monitor cellular morphology and colony-forming efficiency. **d,** Growth curve of untreated and auxin-treated MOF degron mESCs. Data represents mean+/-s.e.m., $n$=3 independent experiments. **e,** Time-course RNA-Seq analyses of MOF degron mESCs ($n$=3 independent experiments) upon auxin treatment. Volcano plots show all expressed transcripts with horizontal dashed line indicating Padj<0.05 significance cut-off and vertical dashed line indicating the log2 fold change cutoff (LFC< -1 & LFC>1). DESeq analysis was performed taking untreated cells as reference for all the conditions. **f,** Pie chart depicting the percentage distribution of H4K16ac peaks with different chromatin features obtained from chromHMM analysis. **g, h,** Example genome snapshots depicting the enrichment of H4K16ac ChIP-Seq signal over promoters and gene bodies **(g)** and enhancers **(h)**. **i,** Schematic depicting the enrichment of BET BRD members BRD2, BRD3 and BRD4 on acetylated chromatin. **j,** Metagene plots showing RPM-normalized ChIP-seq enrichment of BRD2 upon auxin treatment of MOF degron mESCs for 3 h. The signal was plotted over BRD2 ChIP-Seq peaks that were grouped based on their response to MOF depletion shown in **Fig. 1e**. **k,** Metagene plot and heatmap of spike in-normalized native ChIP-Seq signal for H4K5ac, H4K8ac, H4K12ac and H4K16ac over MOF-sensitive BRD2 peaks upon auxin treatment of MOF degron mESCs for 3 h.

Extended Data Fig. 2

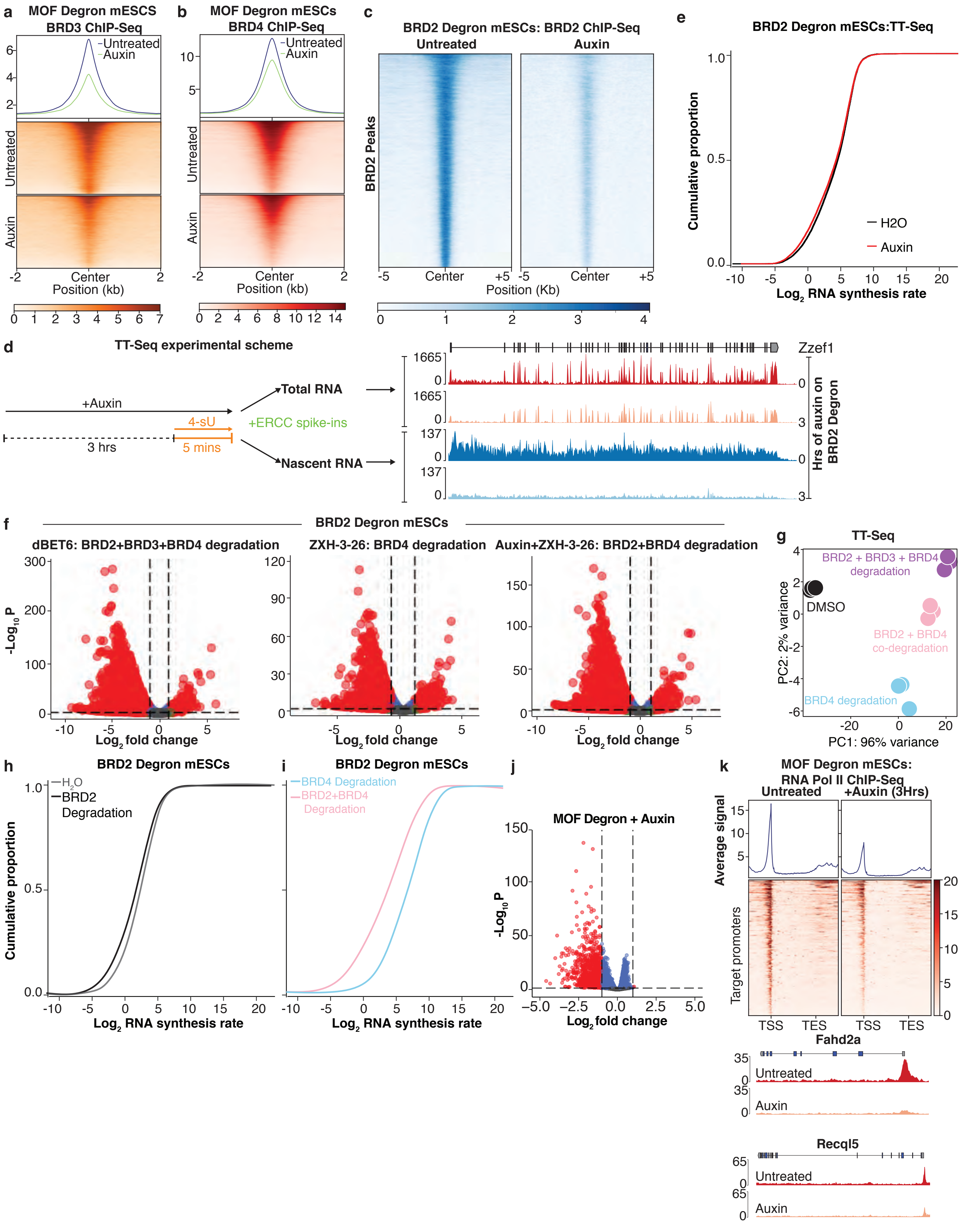

**Extended Data Fig. 2: The distinct roles of BET BRD proteins in promoting nascent RNA synthesis.**

**a, b,** Metagene plot and heatmap showing the ChIP-Seq enrichments of BRD3 **(a)** and BRD4 **(b)** over MOF-sensitive BRD2 peaks upon 3 h auxin treatment of MOF degron mESCs. **c,** Heatmap depicting the input-normalized ChIP-Seq signal of BRD2 upon 3 h auxin treatment of BRD2 degron mESCs. The signal was plotted over BRD2 peaks and the chromatin IP of BRD2 was performed using V5 antibody. **d,** Schematics depicting the experimental strategy for TT-Seq experiments (left). Genome snapshots of total RNA-Seq and TT-Seq signals over an example gene (*Zzef1*) upon auxin treatment of BRD2 degron mESCs (right). **e,** ECDF plot of global RNA synthesis rates from control and BRD2-depleted mESCs. **f,** Volcano plots showing all expressed transcripts with horizontal dashed line indicating Padj<0.05 significance cut-off and vertical dashed line indicating the log2 fold change cutoff (LFC< -1 & LFC>1). DESeq analysis was performed taking DMSO-treated cells as reference for all the conditions. **g,** PCA of TT-seq experiments depicting the clustering of the samples based on their treatment. **h, i,** ECDF plots showing the impact of BRD2 depletion on RNA synthesis rates for the differentially expressed genes from **Fig. 2j** in the presence **(h)** or absence **(i)** of BRD4. **j,** Volcano plot showing differentially expressed genes identified by TT-Seq (red) upon 3-h auxin treatment of MOF degron mESCs. The dashed lines mark the log2FC-cutoff of 1 (vertical lines) and adjusted-p value cutoff of 0.05 (horizontal lines). **k,** Metagene plot, heatmap (top) and example genome snapshots (bottom) of spike in-normalized RNA Pol II ChIP-Seq signal from MOF degron mESCs upon auxin treatment for 3 h. The signal was plotted over the differentially expressed genes obtained from **(j)**.

**Extended Data Fig. 3**

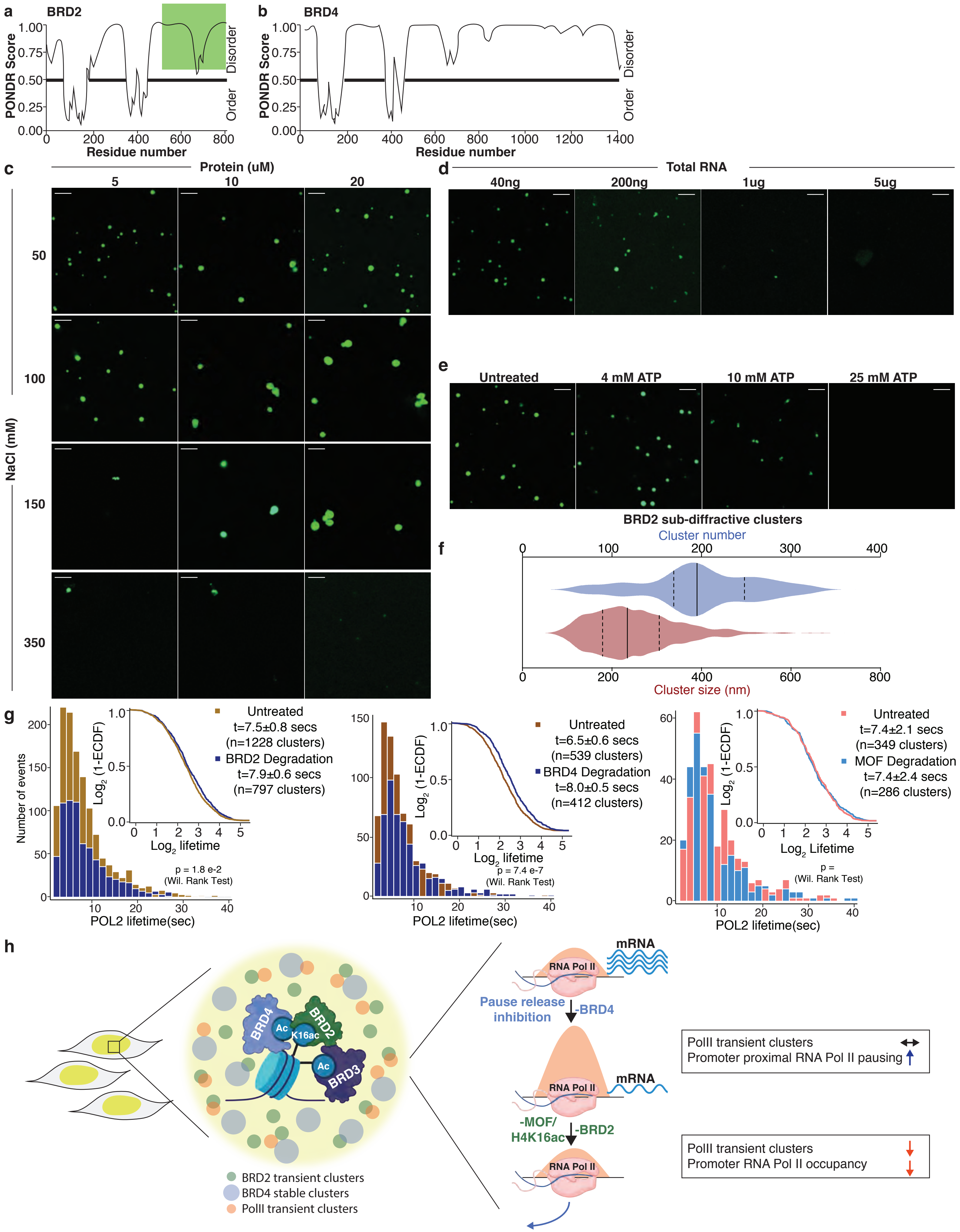

**Extended Data Fig. 3: *In vitro* characterization of BRD2-IDR and working model.**

**a, b,** Graphs plotting the intrinsic disorder score for BRD2 **(a)** and BRD4 **(b)**. PONDR (Predictor of Natural Disordered Regions) VSL2 scores, i.e. neural network predictions of short and long disordered regions, are shown on the y axis and amino acid positions are shown on the x axis. The green box in **(a)** designates the IDR investigated in the lower panels. **c,** Fluorescence microscopy images of the mEGFP-tagged IDR of BRD2 at indicated protein concentrations in buffers with variable NaCl concentrations. Scale bar of images: 5 µm. **d, e,** Fluorescence microscopy images of the purified mEGFP-tagged IDR of BRD2 in buffers with variable total RNA **(d)** or ATP concentrations **(e)**. **f,** Violin plots depicting the BRD2 cluster number (blue) and the cluster size (red) obtained from live-cell super resolution microscopy of BRD2-Halo mESCs. **g,** Histogram and ECDF plots showing the distribution of RNA Pol II cluster lifetimes before and after rapid BRD2, BRD4 and MOF depletion. **h,** A schematic working model depicting the role of BRD2 in the control of paused RNA Pol II dynamics.
